# Dissociating time and number in reinforcement rate learning

**DOI:** 10.1101/302372

**Authors:** Joseph M. Austen, Corran Pickering, Rolf Sprengel, David J. Sanderson

## Abstract

Animals are highly sensitive to the temporal properties of predictive cues ^1^, a feature of learning unexplained by traditional trial-based associative theories of learning ^2-4^. It has been proposed that animals symbolically encode temporal durations in order to extract statistical information about events, such as the reinforcement rate during a cue’s presentation ^5,6^. Thus, long-duration cues elicit weaker responding than short-duration cues (‘cue duration effect’), because they signal a lower rate of reinforcement ^7,8^. An alternative, simple account is that time-dependent changes in stimulus processing, such as short-term habituation, lead to the effect of cue duration on learning ^9,10^, with attention decreasing throughout the duration of a cue, limiting the learning that can occur. Here we provide evidence against the short-term habituation account and show that learning about temporal and numeric information, the quantitative information necessary for rate calculation, can be dissociated by deletion of the GluA1 subunit of the AMPA receptor in mice. In normal mice, the cue duration effect was abolished by equating the cumulative reinforcement rate between cues of different durations across trials. In contrast, GluA1 knockout mice were insensitive to rate information, and their learning was, instead, sensitive to the number of times a cue had been paired with reinforcement over trials. The failure of GluA1 knockout mice to weight learning about the number of reinforcements by the cumulative duration of exposure to a cue was not due to a deficit in short-term habituation or learning about the absence of reinforcement (extinction learning). Instead, it reflected impaired timing ability. The results provide support for models of explicit coding of informational variables achieved through symbolic knowledge ^5,6^, or through time-sensitive learning mechanisms ^11^.

In order to identify the psychological mechanisms underlying sensitivity to temporal properties of cues we began by testing the short-term habituation account in mice with a knockout of *Gria1*, the gene that encodes the GluA1 subunit of the AMPA receptor for glutamate ^12^. GluA1 is necessary for short-term habituation ^13^ and while mice that lack GluA1 (*Gria1*^−/−^ mice) show impairments in short-term habituation ^14-17^ their long-term learning and memory is normal ^12,14,18,19^. Therefore, if the cue duration effect is caused by short-term habituation then GluA1 deletion should reduce the effect and increase learning with a long duration cue.

Mice received Pavlovian conditioning with a short duration, 10 s cue (e.g., light) and a long duration, 40 s cue (e.g., noise) that both terminated with the presentation of reinforcement (sucrose pellet). Normal, wild-type mice showed the cue duration effect, learning more about the short than long cue (Fig 1a). Consistent with the short-term habituation account of the cue duration effect, *Gria1*^−/−^ mice failed to show the cue duration effect and responding between the cues was similar (Fig 1b; cue duration x genotype x block interaction: F_(11,253)_ = 2.66, p = 0.023).

**Figure 1.**
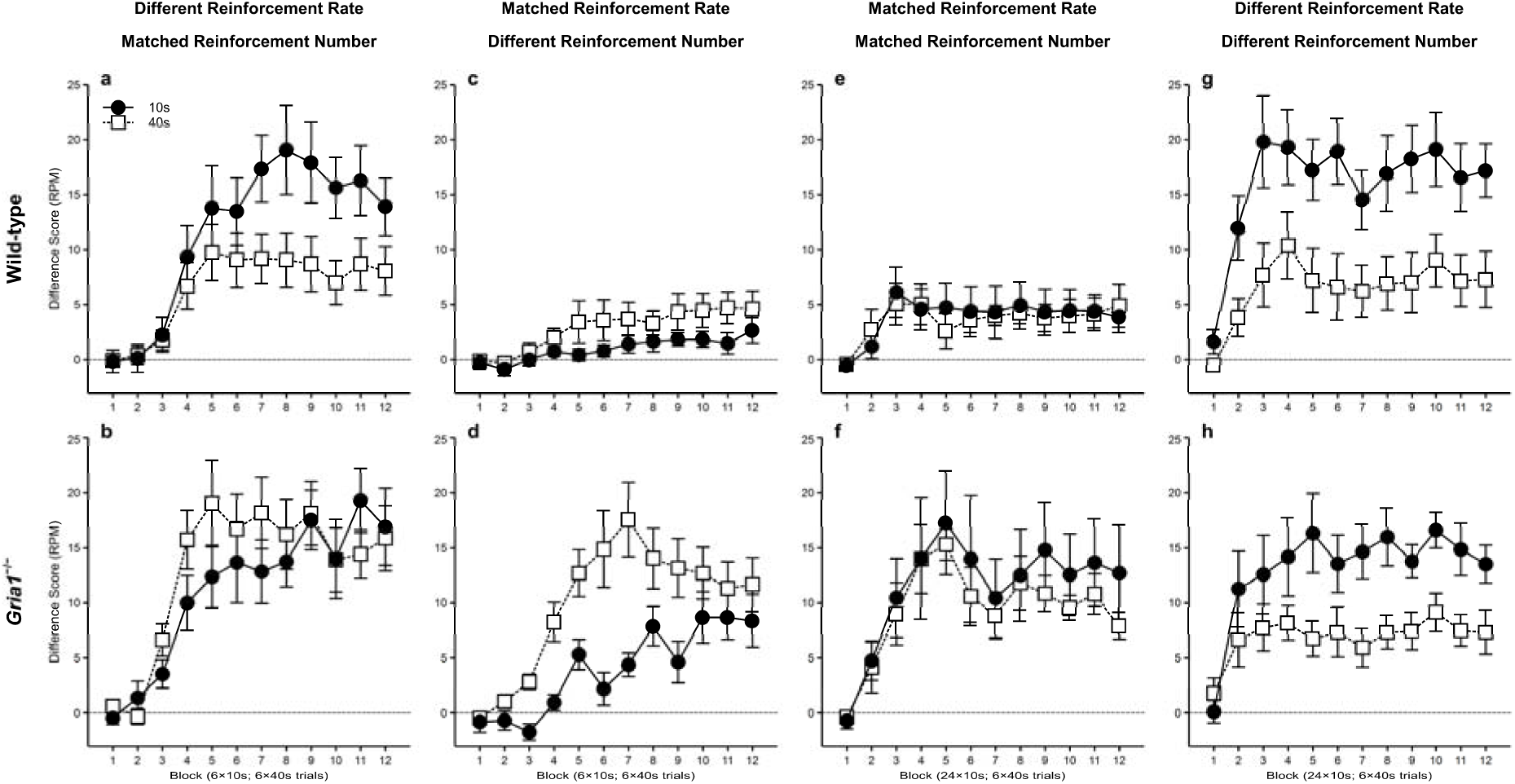
GluA1 deletion switches learning from being dependent on reinforcement rate to being dependent on number of reinforcements. Responding (food magazine entries) is shown minus the response rate to a nonreinforced cue (difference score, RPM) matched for cue duration and frequency of presentation. Panels a and b: cues differed in reinforcement rate, but not number. Panels c and d: cues were matched for reinforcement rate, but not number. Panels e and f: cues were matched for reinforcement rate and number. Panels g and h: cues differed in reinforcement rate and number. Error bars indicate ±SEM.

While the results are consistent with the short-term habituation account of the cue duration effect, it is possible that, instead, GluA1 deletion abolished the cue duration effect by impairing calculation of reinforcement rate. In order to test the role of GluA1 in reinforcement rate learning we trained naïve mice with short and long duration cues, but now the short duration cue was reinforced on one in four trials, such that the cumulative rate of reinforcement across the short and long duration cues was matched (1/40 s). Consistent with reinforcement rate being a key determinant of learning, the cue duration effect was now abolished in wild-type mice, with learning with the short cue being similar to the long-duration cue (Fig 1c). *Gria1*^−/−^ mice, however, now showed superior learning with the long cue than compared to the short cue (Fig 1d; cue duration x genotype x block interaction: F_(11,308)_ = 2.87, p = 0.022). This may suggest that GluA1 deletion impairs sensitivity to reinforcement rate, but an alternative explanation is that the poor learning with the short duration cue was due to the four-fold reduction in the number of pairings with reinforcement rather than the reduction in reinforcement rate. Therefore, in order to test whether the learning with the short cue was affected by reinforcement rate rather than simply reinforcement number we repeated the procedure in naïve mice except that now the partially reinforced short cue was presented four times as often as the long duration cue, such that the short and long duration cues were matched for both reinforcement rate and number of reinforcements. The increase in number of reinforcements failed to increase learning with the short cue compared to the long cue in wild-type mice (Fig 1e), demonstrating that learning in wild-type mice is sensitive to reinforcement rate. In contrast, the increase in the number of reinforcements resulted in learning with the short cue now no longer differing from the long cue in *Gria1*^−/−^ mice (Fig 1f; effect of cue, and cue duration x genotype and cue duration x genotype x block interactions: F-values < 1, p-values > 0.4).

These results suggest that *Gria1*^−/−^ mice are insensitive to reinforcement rate, but are instead sensitive to the number of reinforcements. According to this analysis, *Gria1*^−/−^ mice will show superior learning with the short cue compared to the long cue if the short cue is reinforced more often than the long duration cue. This was tested by training naïve mice with the short and long cues, with each trial reinforced, but the short cue was presented four times as often as the long cue. As predicted, both *Gria1*^−/−^ mice and wild-type mice showed greater learning with the short cue compared to the long cue (Fig 1g-h; effect of cue: F_(1,14)_ = 71.1, p < 0.001; all interactions: F-values < 2.1, p-values > 0.11). Furthermore, because the total cumulative exposure to the short and long duration cues was matched, the results demonstrate that the weaker learning with the long cue compared to the short cue in wild-type mice does not reflect reduced learning caused by the relative familiarity of the cue ^20,21^.

The previous experiments indicate that GluA1 has a role in reinforcement rate calculation over and above its role in short-term habituation. However, reinforcement rate was equated between cues by confounding the probability of reinforcement per trial with the duration of the cue (i.e., a 10 s cue that was reinforced on 25% of trials was compared to a 40 s cue that was reinforced on 100% of trials). Therefore, rate sensitivity may have been achieved by an interaction between the effects of shortterm habituation and partial reinforcement, rather than rate calculation. If GluA1 is important for reinforcement rate calculation then GluA1 will be necessary for sensitivity to reinforcement rate even when cues are of the same duration. We tested this prediction by training naïve mice with two cues that were both 10 s in duration: one cue was reinforced on every trial (high reinforcement rate, 1/10 s), but the other was reinforced on one in every four trials (low reinforcement rate, 1/40 s). The low reinforcement rate cue was presented four times as often as the high reinforcement rate cue such that both cues received the same number of pairings with reinforcement. Wild-type mice showed greater learning with the high reinforcement rate cue than with the low reinforcement rate cue (Fig 2a). Consistent with GluA1 having a role in rate calculation beyond any effect on short-term habituation, *Gria1*^−/−^ mice showed equal learning with the high and low reinforcement rate cues (Fig 2b; cue x genotype interaction: F_(1,20_) = 9.67, p = 0.006).

**Figure 2.**
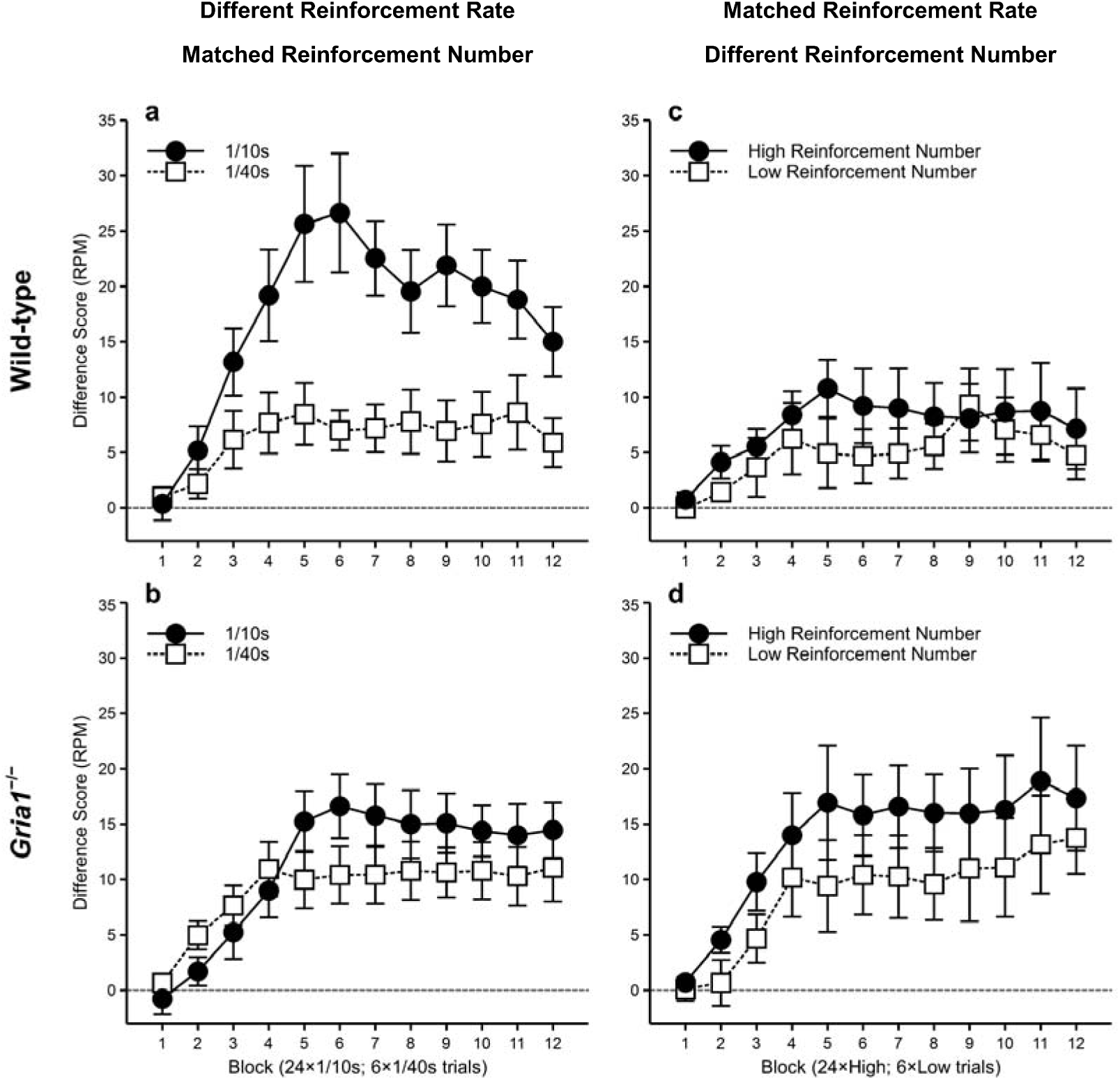
GluA1 deletion impairs sensitivity to reinforcement rate, but not number of reinforcements when cue duration is matched between cues. Responding (food magazine entries) is shown minus the response rate to a nonreinforced cue (difference score, RPM) matched for frequency of presentation. Panels a and b: cues differed in reinforcement rate, but not number. Panels c and d: cues were matched for reinforcement rate, but not number. Error bars indicate ±SEM.

While GluA1 deletion impaired sensitivity to reinforcement rate, it did not impair sensitivity to the number of times a cue was paired with reinforcement. Indeed, *Gria1*^−/−^ mice were sensitive to reinforcement number under conditions in which wild-type mice were not (Fig 1c-d). In order to test directly sensitivity to number of reinforcements naïve mice were tested on a procedure in which cue duration and reinforcement rate were matched; two 10 s cues were both reinforced one in every four trials (reinforcement rate = 1/40s), but one cue (high reinforcement number) was presented four times more often than the other cue (low reinforcement number). Overall, independent of genotype, mice showed superior learning with the high reinforcement number cue than with the low reinforcement number cue (Fig 2c-d; effect of cue: F_(1,20)_ = 5.57, p = 0.029; all interactions: F-values < 1.6, p-values > 0.19), demonstrating that GluA1 deletion spares sensitivity to reinforcement number.

The results demonstrate that learning in normal mice is sensitive to the rate of reinforcement, regardless of whether reinforcement rate is manipulated by the duration of the cue or the probability of reinforcement per trial. Deletion of GluA1 qualitatively changed learning from being dependent on reinforcement rate to being dependent on the number of pairings with reinforcement, revealing that GluA1 is necessary for weighting numeric information by temporal information for the calculation of reinforcement rate. Sensitivity to temporal information in the predictive learning procedures reflected extraction of statistical information from events. While this may reflect encoding of numeric and temporal variables ^5,6^, it may also be achieved by the balance of increments and decrements in associative strength across cumulative exposure to cues ^11,22,23^. Thus, an error correction rule implemented in an iterative manner over time achieves rate calculation by reducing associative strength during periods of nonreinforcement (see Supplementary Material). This model predicts that GluA1 deletion impairs the decrements in learning during nonreinforcement (i.e., learning through negative prediction error ^4,24^). We have, however, failed to find support for this account, with *Gria1*^−/−^ mice showing normal extinction of learning during nonreinforced exposure to a previously reinforced cue (Fig 3a-b; cue x block interaction: F_(7,154)_ = 11.68, p < 0.001; cue x genotype and cue x genotype x block interactions: F-values < 2.6, p-values > 0.12). In contrast, *Gria1*^−/−^ mice failed to time conditioned responses appropriately in terms of linear gradients (Fig 4a; effect of genotype: F_(1,35)_ = 6.90, p = 0.013) or spread of distribution (Fig 4b; effect of genotype: U = 62.0, p = 0.001), a property of predictive learning indicative of memory for the time of reinforcement ^25,26^. Therefore, these results suggest GluA1 deletion impaired rate calculation by impairing the encoding of the representations of temporal durations ^6^.

**Figure 3.**
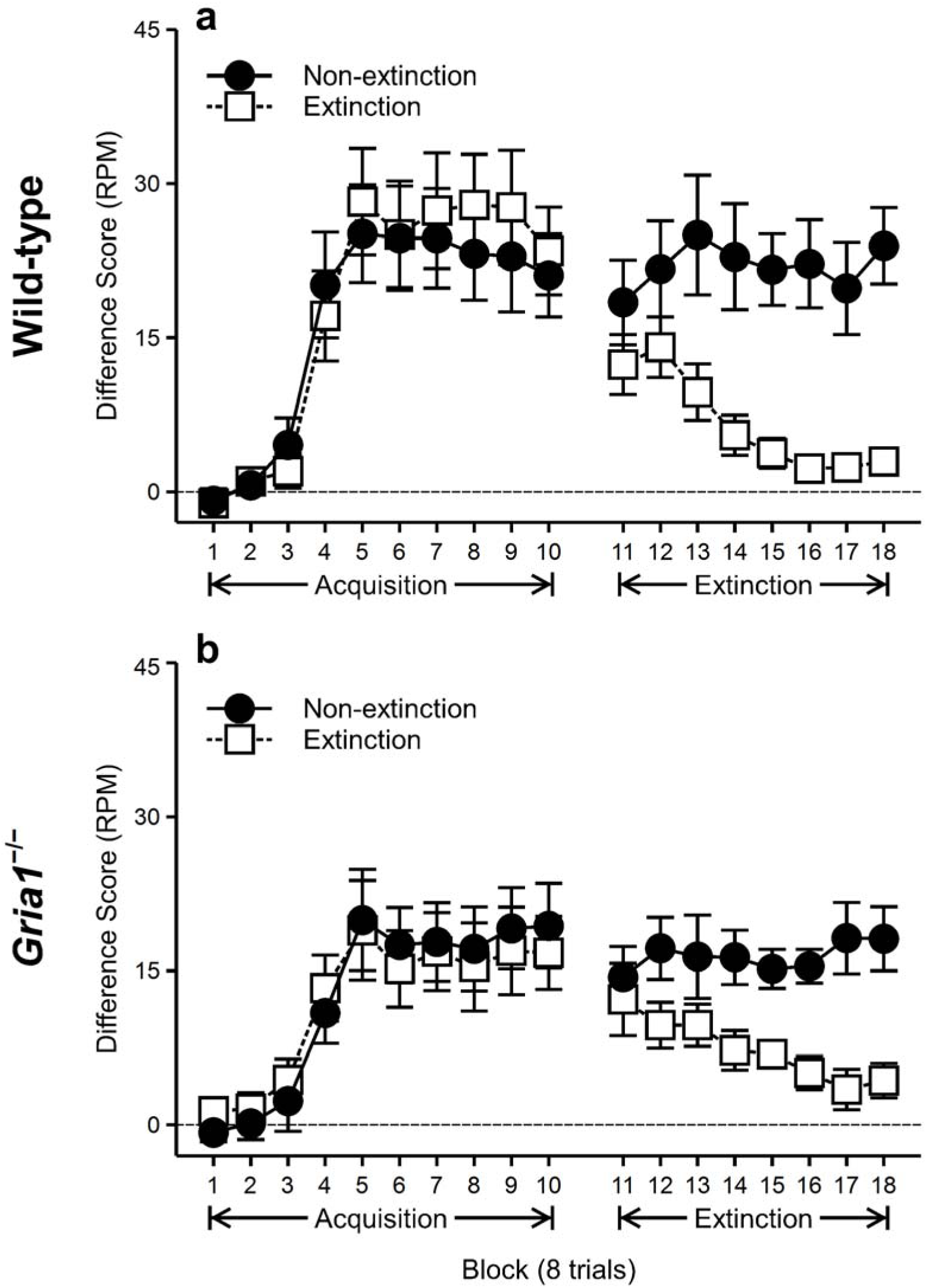
GluA1 deletion does not impair extinction of conditioned responding. Responding (food magazine entries) is shown minus the response rate to a nonreinforced cue (difference score, RPM). Panels a and b: during the acquisition phase (blocks 1-10) mice received training with two 10 s cues that were reinforced on every trial. During the extinction phase (blocks 11-18) the extinction cue was no longer reinforced, but the non-extinction cue continued to be reinforced. Error bars indicate ±SEM.

**Figure 4.**
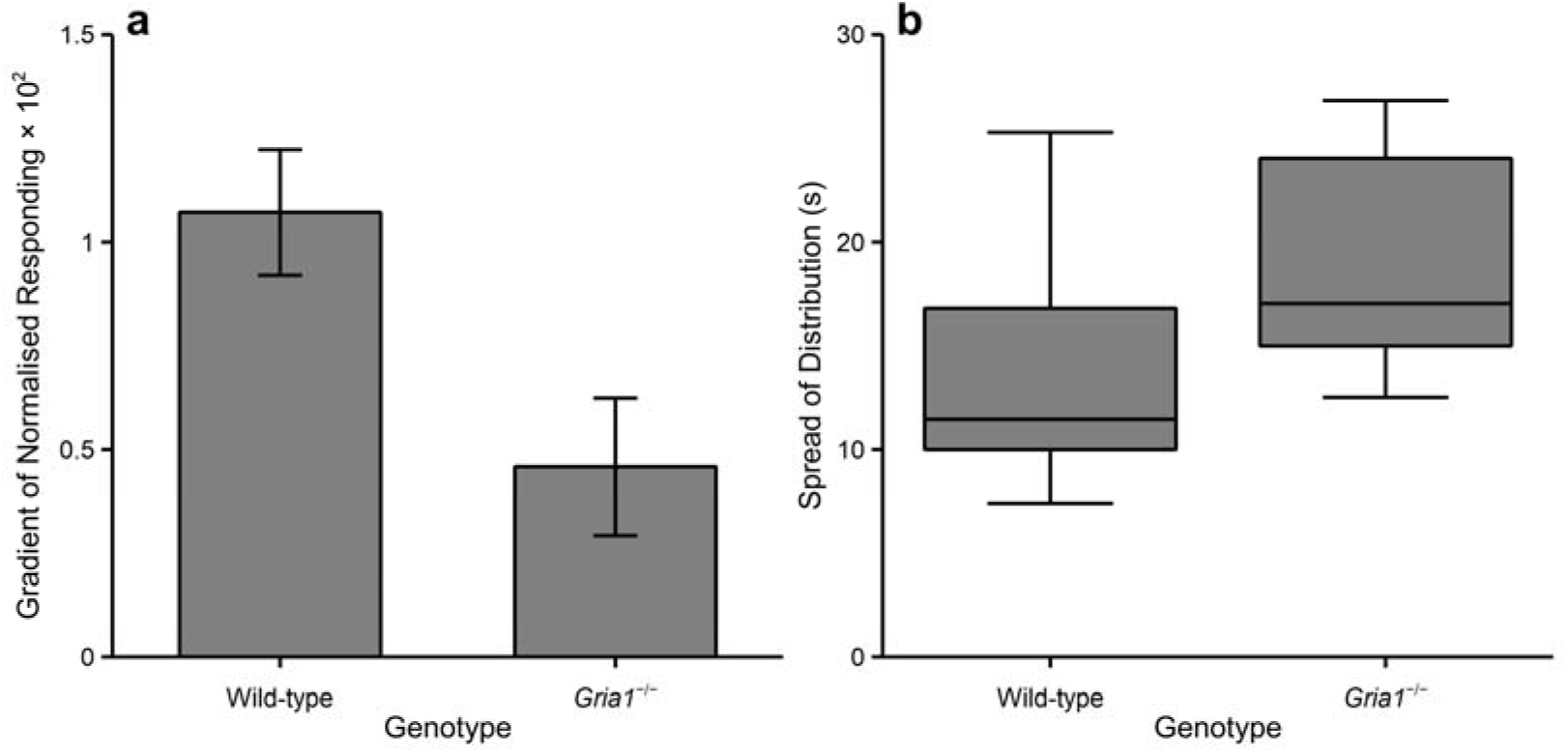
GluA1 deletion impairs timing of conditioned responses. (a) GluA1 deletion reduced the gradient of the linear slopes fitted to the first 10 s of the 30 s cue (prior to reward delivery). Error bars indicate ±SEM. (b) GluA1 deletion increased the spread of the timing distribution modelled with a Gaussian function. Whiskers represent 1.5IQR.

Some traditional trial-based associative learning models have appealed to the temporal dynamics of stimulus processing, such as memory decay, in order to explain temporal sensitivity in learning ^4,27^. Here, we used a novel test of a time-dependent processing account: genetic manipulation of short-term habituation. While GluA1 deletion did affect learning, it was clear that the role of GluA1 was not in determining time-dependent changes in stimulus processing, but was actually in calculation of statistical information about the environment. These results provide new evidence that learning reflects the encoding of the details and structure of events through either symbolic knowledge ^5,28,29^, or as an emergent property of time-sensitive learning processes ^11^.

## Methods

### Subjects

Mice were *Gria1*^−/−^ and wild-type age-matched littermates, bred in the Life Sciences Support Unit, Durham University (see Zamanillo et al. (1999) for details of genetic construction, breeding and subsequent genotyping). The mice were originally derived from the 129S2svHsd and C57BL/6J/OlaHsd strains, and have been subsequently backcrossed onto the C57BL/6J line. Mice were housed in groups of 1-12 in a temperature controlled holding room on a 12-hour light-dark cycle (light period: 8am to 8pm). For several days prior to the start of testing, the weights of the mice were reduced by restricting access to food and they were maintained at 85% of their free-feeding weights throughout the experiment. Mice had ad libitum access to water in their home cages. All procedures were in accordance with the United Kingdom Animals Scientific Procedures Act (1986); under project license number PPL 70/7785.

### Apparatus

A set of eight identical operant chambers (interior dimensions: 15.9 × 14.0 × 12.7 cm; ENV-307A, Med Associates), enclosed in sound-attenuating cubicles (ENV-022V) were used. The operant chambers were controlled by Med-PC IV software (SOF-735). The side walls were made from aluminium, and the front and back walls and the ceiling were made from clear Perspex. The chamber floors each comprised a grid of stainless steel rods (0.32 cm diameter), spaced 0.79 cm apart, running perpendicular to the front of the chamber (ENV-307A-GFW). A food magazine (2.9 × 2.5 × 1.9 cm; ENV-303M) was situated in the centre of one of the sidewalls of the chamber, into which sucrose pellets (14 mg, TestDiet) could be delivered from a pellet dispenser (ENV-203-14P). An infrared beam (ENV-303HDA) across the entrance of the magazine recorded head entries at a resolution of 0.1 s. A fan (ENV-025F) was located within each of the sound-attenuating cubicles and was turned on during sessions, providing a background sound level of approximately 65 dB. Auditory stimuli were provided by a white noise generator (ENV-325SM) that outputted a flat frequency response from 10 to 25,000 Hz at 80 dB, a clicker (ENV-335M) that operated at a frequency of 4 Hz at 80 dB, and a pure tone generator (ENV-323AM) that produced a 2,900 Hz tone at 80dB. Visual stimuli were a 2.8 W house light (ENV-315M), and two LEDs (ENV-321M) that were flashed (1 s on/1 s off) alternating between left and right.

### Cue duration experiments

#### General procedures

Mice received training, one session per day, with two 10 s cues and two 40 s cues. One cue of each duration was reinforced (either on 100% or 25% of those trials depending on the experiment) by presentation of a sucrose pellet at the termination of the cue. The remaining cues were not reinforced. Trials were separated by a fixed interval of 120 s (cue offset to cue onset). For approximately half of the mice within each genotype and sex the 10 s cues were visual (house light, LEDs) and the 40 s cues were auditory (white noise, clicker). Within modality the allocation of reinforced and nonreinforced cues was counterbalanced as far as possible. For the remaining mice the 10 s cues were auditory and the 40 s cues were visual, and the reinforcement contingencies within modality were similarly counterbalanced.

#### Different reinforcement rate – matched reinforcement number

Thirteen *Gria1*^−/−^ (7 female, 6 male) and 14 wild-type (6 female, 8 male) mice (free-feeding weights: 14.6 – 28.1 g) received 12 sessions of training in which the reinforced 10 s and 40 s cues were reinforced on every trial. Sessions consisted of six trials of each trial type (10 s reinforced, 10 s nonreinforced, 40 s reinforced and 40 s nonreinforced). The cues were presented in a random order with the constraint that there was an equal number of each cue type every eight trials.

#### Matched reinforcement rate – different reinforcement number

Sixteen *Gria1*^−/−^ (9 female, 7 male) and 16 wild-type (9 female, 7 male) mice (free-feeding weights: 16.2 – 33.2 g) received 12 sessions of training in which the 10 s reinforced cue was reinforced on only 25% of trials and the 40 s reinforced cue was reinforced on every trial. Sessions consisted of six trials of each trial type (10 s reinforced, 10 s nonreinforced, 40 s reinforced and 40 s nonreinforced). The partially reinforced 10 s cue was reinforced either once or twice per session, and it was ensured that across consecutive pairs of sessions the 10 s cue was reinforced three times. The cues were presented in a random order with the constraint that there was an equal number of each cue type every eight trials.

#### Matched reinforcement rate – matched reinforcement number

Seven *Gria1*^−/−^ (3 female, 4 male) and 10 wild-type (8 female, 2 male) mice (free-feeding weights: 16.4 – 29.7 g) received 24 sessions of training in which the 10 s reinforced cue was reinforced on only 25% of trials and the 40 s reinforced cue was reinforced on every trial. Sessions consisted of 12 trials of each of the 10 s cues (reinforced and nonreinforced) and 3 trials of each of the 40 s cues (reinforced and nonreinforced). The cues were presented in a random order with the constraint that there were four trials of each 10 s cue and 1 trial of each 40 s cue every 10 trials. Therefore, every 10 trials the reinforced 10 s cue was reinforced once.

#### Different reinforcement rate – Different reinforcement number

Eight *Gria1*^−/−^ (3 female, 5 male) and 10 wild-type (7 female, 3 male) mice (free-feeding weights: 15.9 – 33.0 g) received 24 sessions of training in which the 10 s and 40 s reinforced cues were reinforced on every trial. Sessions consisted of 12 trials of each of the 10 s cues (reinforced and nonreinforced) and 3 trials of each of the 40 s cues (reinforced and nonreinforced). The cues were presented in a random order with the constraint that there were four trials of each 10 s cue and 1 trial of each 40 s cue every 10 trials.

### Matched cue duration experiments

#### Different reinforcement rate – matched reinforcement number

Thirteen *Gria1*^−/−^ (7 female, 6 male) and 11 wild-type (5 female, 6 male) mice (free-feeding weights: 18.2 – 34.1 g) received 24 sessions of training, one per day, in which two 10 s cues (12 trials each per session) were presented four times as often as two other 10 s cues (3 trials each per session). Trials were separated by a fixed interval of 120 s (cue offset to cue onset).One of the more frequently presented cues was reinforced (by presentation of a sucrose pellet at the termination of the cue) on 25% of trials (reinforcement rate = 1/40 s) and the other was nonreinforced. One of the less frequently presented cues was reinforced on every trial (reinforcement rate = 1/10 s) and the other was nonreinforced. For approximately half of the mice within each genotype and sex the more frequently presented cues were visual (house light, LEDs) and the less frequently presented cues were auditory (white noise, clicker). Within modality the allocation of reinforced and nonreinforced cues was counterbalanced as far as possible. For the remaining mice the more frequently presented cues were auditory and the less frequently presented cues were visual, and the reinforcement contingencies within modality were similarly counterbalanced. The cues were presented in a random order with the constraint that there were four trials of each of the more frequently presented 10 s cues and 1 trial of each of the less frequently presented cues every 10 trials. Therefore, every 10 trials the reinforced 10 s cue was reinforced once.

#### Matched reinforcement rate – different reinforcement number

Twelve *Gria1*^−/−^ (6 female, 6 male) and 12 wild-type (4 female, 8 male) mice (free-feeding weights: 17.3 – 33.8 g) received training that was the same as the *Different rate – matched reinforcement number* experiment when cue durations were matched, except that the less frequently presented reinforced cue was reinforced on only 25% of trials.

### Extinction

Twelve *Gria1*^−/−^ (6 female, 6 male) and 12 wild-type (6 female, 6 male) mice (free-feeding weights: 17.4 – 33.4 g) initially received 10 sessions of training, one session per day, in which three 10 s cues were presented eight times each per session. Trials were separated by a fixed interval of 120 s (cue offset to cue onset). Two of the cues were reinforced (by presentation of a sucrose pellet at the termination of the cue) on every trial and one cue was nonreinforced. The order of trials was random with the constraint that there was an equal number of each trial type every six trials. In the second stage, which lasted for eight sessions, all procedures were the same except that one previously reinforced cues was now nonreinforced (extinction). The other previously reinforced cue continued to be reinforced (non-extinction). The three stimuli were white noise, tone and clicker. The allocation of stimuli to trial types was counterbalanced within genotype and sex.

### Timing

Fourteen *Gria1*^−/−^ (7 female, 7 male) and 23 wild-type (12 female, 11 male) mice (free-feeding weights: 20.9 – 32.6 g) received 12 sessions of training, one session per day, in which two 30 s cues were presented twelve times each per session. Trials were separated by a fixed interval of 120 s (cue offset to cue onset). One of the cues was reinforced (by presentation of a sucrose pellet 10 s after cue onset) on 75% of trials and the other cue was nonreinforced. The order of trials was random with the constraint that there was an equal number of each trial type every eight trials, with the reinforced cue being rewarded on three of its four trials. The two stimuli were white noise and clicker. The allocation of stimuli to trials types was approximately counterbalanced within genotype and sex.

### Data and statistical analysis

For all experiments, the number of head entries made to the food magazine during the presentation of each CS was recorded. Responding during nonreinforced cues was then subtracted from responding during reinforced cues that were matched for cue duration and/or frequency of presentation, to give levels of responding relative to baseline for each condition. All data, except the timing experiment (see below) were analysed using multifactorial ANOVA. For all experiments in which both visual and auditory cues were used, counterbalancing of modality was included as a nuisance factor (due to mice responding more to auditory than visual cues). Interactions were analysed with simple main effects analysis using the pooled error term from the original ANOVA, or separate repeated measures ANOVA for within-subject factors with more than two levels. Where sphericity of within-subjects variables could not be assumed, a Greenhouse-Geisser correction was applied to produce more conservative *p*-values. The key main effects and interactions are reported in the main text; the full analyses are reported in the Supplementary Material.

#### Analysis of timing of conditioned responding

Timing of responding was assessed in two ways: linear slopes and Gaussian curve-fitting. Linear slopes: Responding during the first 10 s of the reinforced cue (i.e., prior to the delivery of reward) was divided into ten equal time periods (i.e., ten one-second time bins) and averaged across all sessions of the experiment. These data were then normalised to show the proportion of responding an animal made during each of the ten time bins. The linear gradients of these normalised data were then calculated to provide an indication of the extent that responding to cues was being timed (i.e., the steeper the gradient the more the animals were timing their responding to the delivery of the US).

Gaussian curve-fitting: Responding during each 1 s time bin of the entire 30 s of the non-rewarded presentations of the reinforced cue was averaged across all trials and sessions and then smoothed over four 1 s bins. A Gaussian model (Eq. 1) was then fitted to the timing distribution of each animal. Here, *R_i_*, is the conditioned responding in smoothed time bin *i*, *A* is the peak rate of responding, *B* is the spread of the distribution, *C* is the time bin at which the peak rate occurs (the central tendency, peak time), and *x_i_* is the time since CS onset in smoothed time bin *i.*

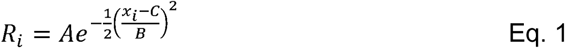

The different measures of timing were analysed using Mann Whitney U tests. The results for the peak rate, peak time, and spread measures, in addition to measures of peak error and coefficient of variation (spread/peak time), are reported in the Supplementary Material.

## Acknowledgements

The work was supported by a grant from BBSRC (BB/M009440/1) to DJS. We thank Young Ah Kim and Alex Finniss for support with behavioural testing.

## Author contributions

The experiments were designed by JMA and DJS. JMA and CP conducted the experiments and analysed the results. JMA, RS and DJS wrote the paper.

